# MicroRNA-145 enhances lung cancer cell progression after exposure to lyophilized fertile hydatid cyst fluid of *Echinococcus granulosus* sensu stricto

**DOI:** 10.1101/2024.01.26.577517

**Authors:** Hosein Mosajakhah, Dariush Shanehbandi, Ehsan Ahmadpour, Mahmoud Mahami-Oskouei, Khadijeh Sadeghi Janghoor, Adel Spotin

**Affiliations:** Immunology Research Center, Tabriz University of Medical Sciences, Tabriz, Iran; Department of Parasitology and Mycology, Faculty of Medicine, Tabriz University of Medical Sciences, Tabriz, Iran

**Keywords:** Hydatid cyst fluid, Lung cancer, MicroRNA-145, Apoptosis, Migration

## Abstract

There is increasing evidence that the secretory/excretory antigens of the larval stage of *Echinococcus granulosus* (hydatid cyst fluid; HCF) can induce both anticancer and oncogenesis effects between parasite-derived metabolites and various cancer lines. The dual role of miR-145 as a tumor suppressor or oncogene has been previously reported in cancers. Nevertheless, it remains unknown, how miR-145 induces apoptosis in HCF-treated lung cancer cells. The fertile HCF was obtained from sheep and subjected to purification and lyophilization. H1299 human lung cancer cells were cultured into two groups: HCF-treated H1299 lung cancer cells and control cells. To evaluate the effects of HCF on the H1299 cells, cell viability was performed by MTT assay. The caspase-3 activity was assessed using fluorometric assay. Furthermore, the mRNA expression of VGEF, Vimentin, caspase-3, miRNA-145, Bax and Bcl-2 genes were characterized by Real-time PCR. A scratch test was done to assess the effects of HCF on cell mobility. MTT assay revealed that there is an increasing slope in the growth of H1299 cells when treated with 60 μg/mL of fertile HCF for 24 h. The fold change of caspase-3, miRNA-145, Bax/Bcl-2 ratio and caspase-3 activity was lower in the HCF-treated H1299 cells than in the control cell. The fold change of VGEF and Vimentin genes in the HCF-treated H1299 cells was higher than that in the control cell. The scratch outcome demonstrated that the mobility of H1299 cells was increased at 24 and 48 hours of scratched time after exposure to HCF. Our results suggest that induction of low expression of miR-145 in patients with hydatid cysts could be a possible oncogenic regulator of lung cancer growth. We conclude that miR-145 may be a promising marker for the diagnosis of lung cancer patients co-infected with pulmonary hydatid cysts. To validate this assumption, further study is needed to assess microRNA profile and potent oncogenes *in vivo* setting.

## 1. Introduction

Cystic echinococcosis (CE) is an important cyclo-zoonotic disease caused by the larval stage of the *Echinococcus granulosus* sensu lato complex (G1-G10) [1,2]. Approximately, 40 million people are at risk among them four million people are infected [1,3]. The hydatid cyst fluid (HCF) of *E. granulosus* is composed of a mixture of glycolipid, antigens, sulfur, carbohydrates, glycoprotein and cyclophilin [4].

Currently, there is rising evidence that the excretory/secretory (metabolites) antigens of the larval stage of *E. granulosus* (hydatid cyst fluid; HCF) can induce both anticancer and oncogenesis effects between parasite-derived metabolites and various cancer lines (Breast cancer, colon cancer and melanoma, etc.,) [5-9]. Nevertheless, the majority of hydatid cysts are localized in liver and lung of humans (as accidental intermediate hosts), the immunological interaction between HCF antigens and lung cancer line has not been fully understood yet[10]. Some studies have shown that antigenic similarities between some helminths and cancer cells may be effective in inhibiting cancer metastasis [11,12].

Lung cancer is one of the most prevalent cancers worldwide with 2.2 million new cases in 2020 [13]. However, very little is known about the potential role of miRNA in the regulation of *Echinococcus* metabolites to influence lung cancer cells [14].

Several studies describe a dual role of miR-145 as a tumor suppressor and oncogene in cancers. However, the triggering function of miR-145 in the initiation of apoptosis in HCF-treated lung cancer cells has not been fully understood. The principal purpose of this study was to evaluate the different concentrations of lyophilized HCF on H1299 human lung cancer cells to understand whether miR-145 can trigger apoptosis and migration of cancer cells after exposure to HCF of *E. granulosus* s.s.

## 2. Materials and methods

### 2.1. Collection, purification of HCF and genotyping

Fertile liver hydatid cysts were obtained from sheep slaughtered in Maragheh and Tabriz slaughterhouses, northwest Iran. Aspirated HCF was centrifuged (800 g for 15 min) to deposit protoscoleces (PSCs). Viability of the PSCs was confirmed by eosin (0.1%) and trypan blue dyes under a light microscope. In order to purify the fertile hydatid cyst fluid, 100 mL of HCF was dialyzed overnight against 5 mM of sodium acetate buffer (0.005 M; pH 5) at 4°C [15]. Bradford protein assay was used to determine its concentration [16].

The genomic DNA of aspirated PSCs was extracted with a commercial kit (Yekta Tajhiz Azma, Iran). The Cytochrome C oxidase subunit 1 mitochondrial gene (Cox1; 444 bp) was amplified by PCR [17]. To show taxonomic status of *Echinococcus* sp. isolate, PCR product was sequenced (Codon Company, Tehran, Iran), edited and characterized by phylogenetic tree.

### 2.2. Lyophilization

Antigens of HCF were lyophilized according to the standard methods after purification [16].

### 2.3. Cell culture and grouping

The H1299 human lung cancer cell was provided from Pasteur Institute of Iran. H1299 cells were cultured in RPMI1640 supplemented with 10% fetal bovine serum (FBS) and 1% double-antibiotics combination. The 5×10^6^ cells/mL were seeded into a 24-well plate. H1299 cells were cultured into two groups including; various concentrations (20 to 80 μg/mL) of HCF (lyophilized HCF) against H1299 cells and control cells. Normal lung epithelial cells were considered as control cells.

### 2.4. MTT assay

The growth properties of HCF on H1299 cells were characterized by MTT (3-(4,5-dimethylthiazol-2yl)-2,5-diphenyltetrazolium bromide) assay after 24 h exposure time. The protocol of MTT assay was previously described by Mohammadi et al [5]. Presence of viable cells was visualized by the development of purple color due to formation of formazan crystals. The plates were measured by using an ELISA reader at 570 nm.

### 2.5. Wound healing assay (Scratch test)

To assess the growth effects of HCF on the migration of H1299 cells, 5 × 10^5^ cells were seeded in six-well cell culture plate in RPMI 1640 medium. After 24 h of incubation, cells filled the entire bottom of wells subsequently; scratch was performed by a sterile yellow sampler tip to each well. Images were taken from each well by an inverted microscope at 0, 24 and 48 h. The mobility of cells was compared between HCF-treated H1299 cells and control cells.

### 2.6. Extraction of RNA, cDNA synthesis and real-time PCR (RT-PCR)

Total RNAs (mRNA and miRNA) were extracted from cells using TRIzol reagent. Complementary DNA (cDNA) was synthesized and harvested from fertile HCF-treated H1299 cells and control cells. For quantitative assessment of Bax, Bcl-2, caspase-3, miR-145, VGEF and Vimentin genes, qRT-PCR was performed. Primers used in this study are given in Table 1. RT-PCR reactions were repeated in triplicate by using GAPDH and U6 as internal controls. To calculate fold changes in expression levels of caspase-3, VGEF, Vimentin, Bax, Bcl-2 and miRNA-145, the Ct (2^−ΔΔCt^) formula was used.

**Table 1:**
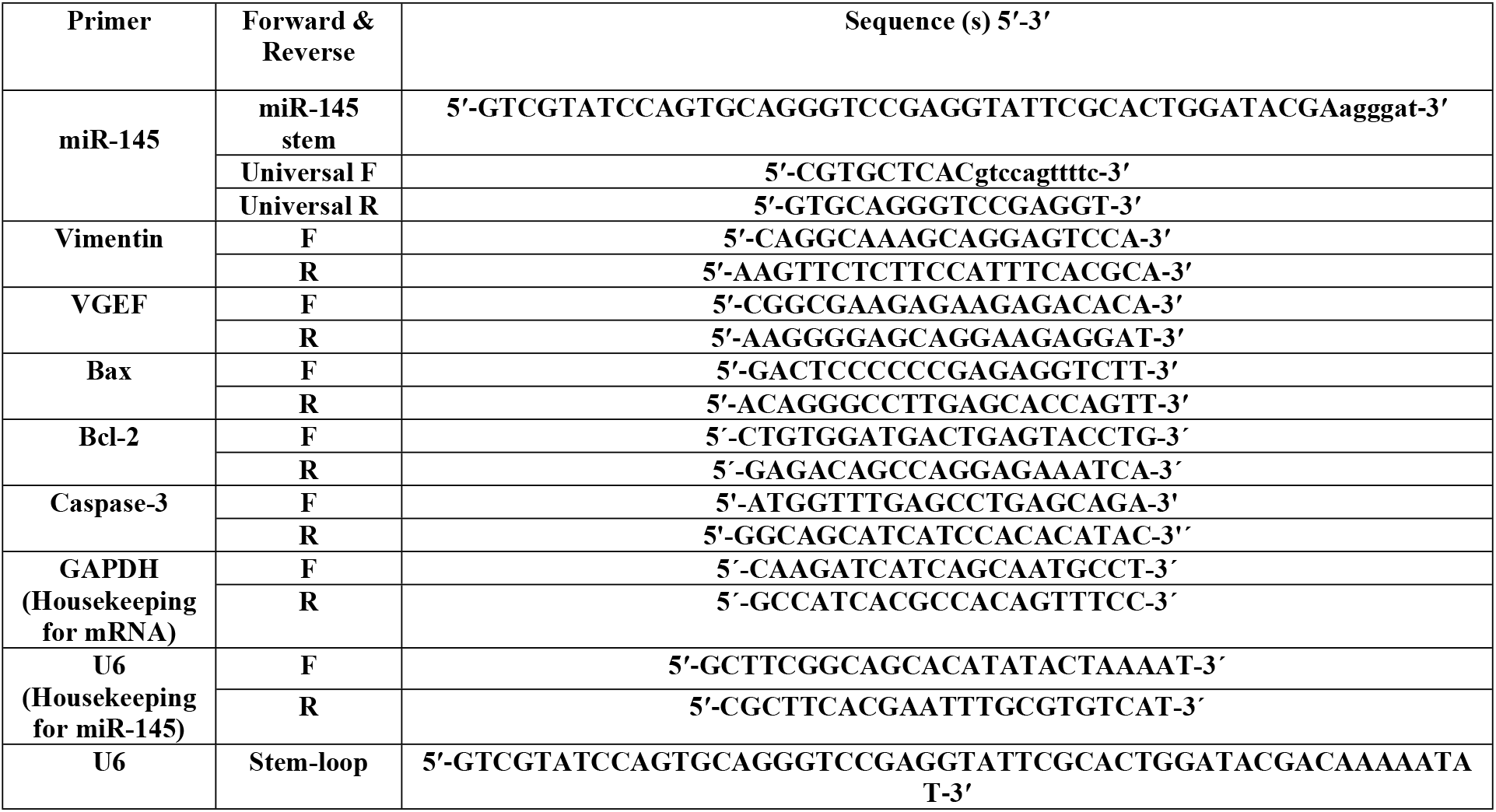
The sequence of primers used in qRT-PCR.

### 2.7. Assessment of caspase-3 activity

The caspase-3 activity was evaluated by the caspase-3/CPP32 Fluorometric Assay Kit after 15 h of exposure to HCF on H1299 lung cancer cell line.

### 2.8. Statistical analysis

The fold change in mRNA expression of assessed genes was analyzed by Wilcoxon signed rank test using GraphPad-Prism-5 software.

## 3. Results

### 3.1. Genotyping and the effects of HCF on H1299 cells

Based on sequencing analysis of *Cox1* gene, sheep strain (G1 genotype) belonging to *E. granulosus* sensu stricto complex (G1-G3) was characterized. *In vitro* MTT assay indicated an increasing slope in the growth of H1299 human lung cancer cells when treated with rising concentrations of HCF (20, 30, 40, 50 and especially 60 μg/mL) during 24 h exposure time (Figure 1).

**Figure 1:**
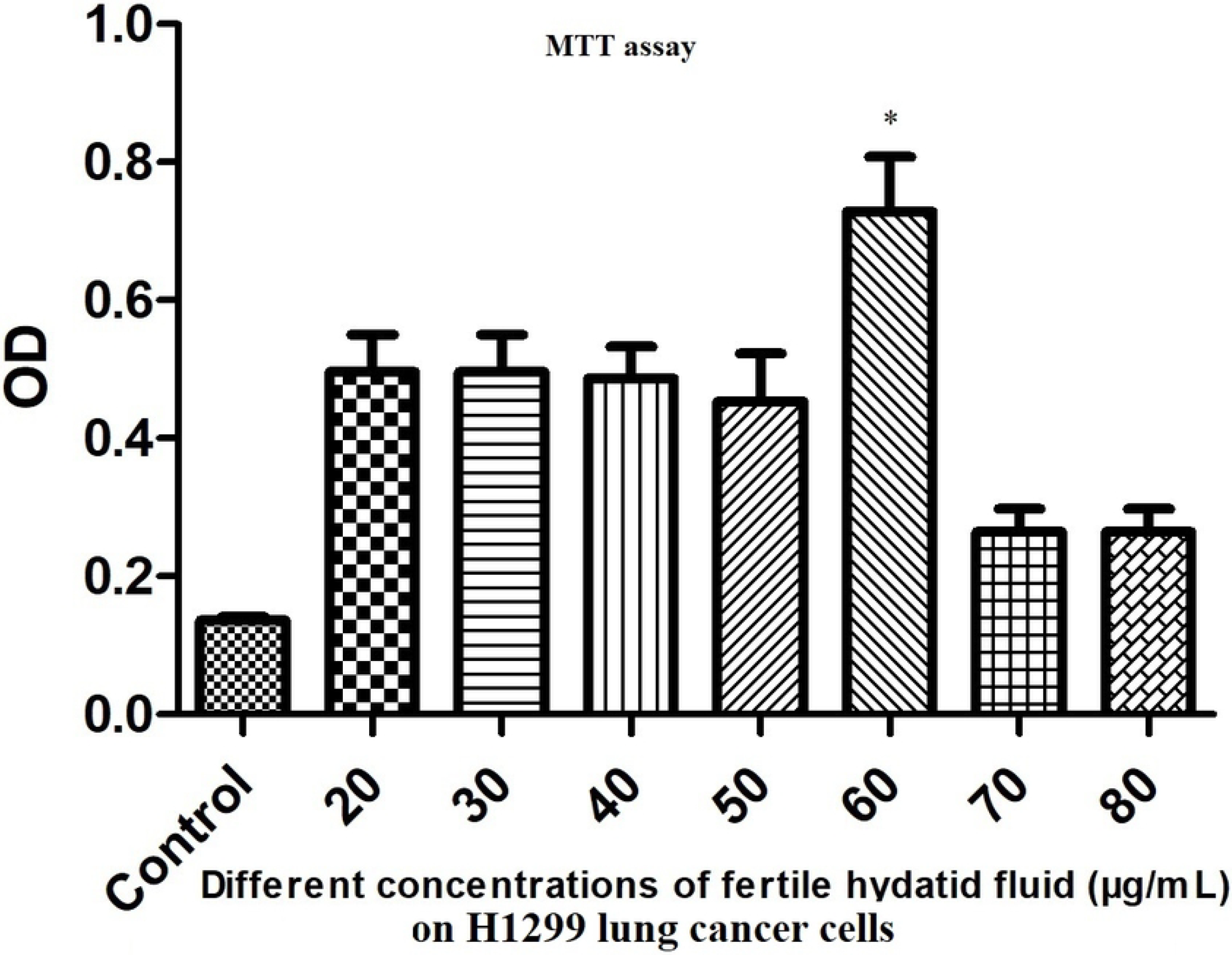
The growth effects of various concentrations of fertile HCF-treated H1299 human lung cancer cells at 570 nm by using a MTT assay after 24 h exposure time.

### 3.2. Assessment of mRNA expression of miR-145

We showed that 70 μg/mL of fertile HCF-treated H1299 cells down-regulated the miR-145 expression level about 2-fold change decrease compared to control cells (Figure 2; *P*v; 0.0021).

**Figure 2:**
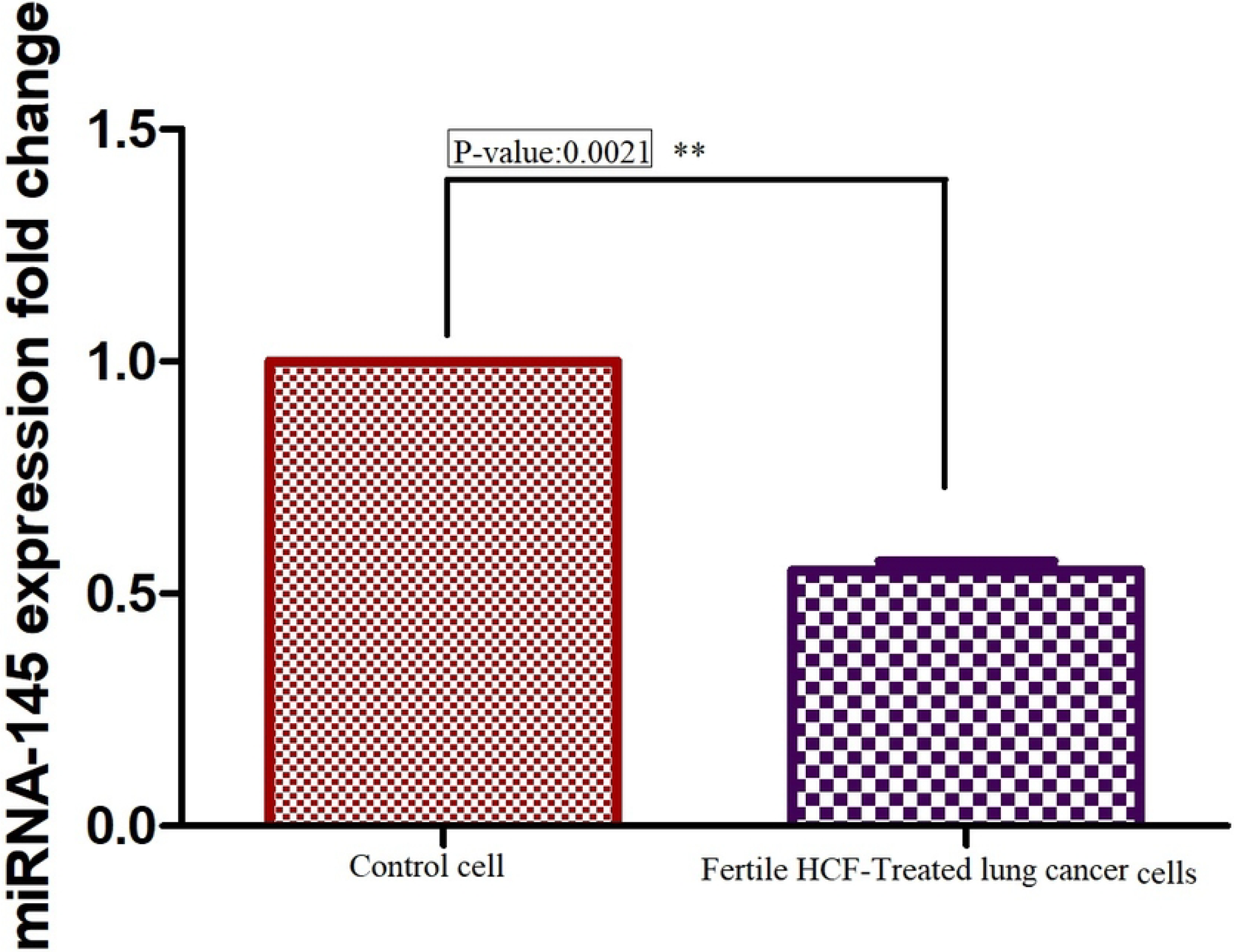
Expression of miR-145 gene in fertile HCF-treated H1299 human lung cancer cells compared to control cells.

### 3.3. Effects of fertile HCF on the migration of H1299 cell line

The scratch test was done to assess the effect of fertile HCF on the migration of the H1299 lung cancer cells. Current findings denoted that the migration of H1299 cells was increased after 24 (188 cells per field to 222 cells per field: *P*v <0.05) and 48 h (156 cells per field to 310 cells per field: *P*v < 0.001) of scratched time following exposure to fertile HCF compared to control cells (Figure 3).

**Figure 3:**
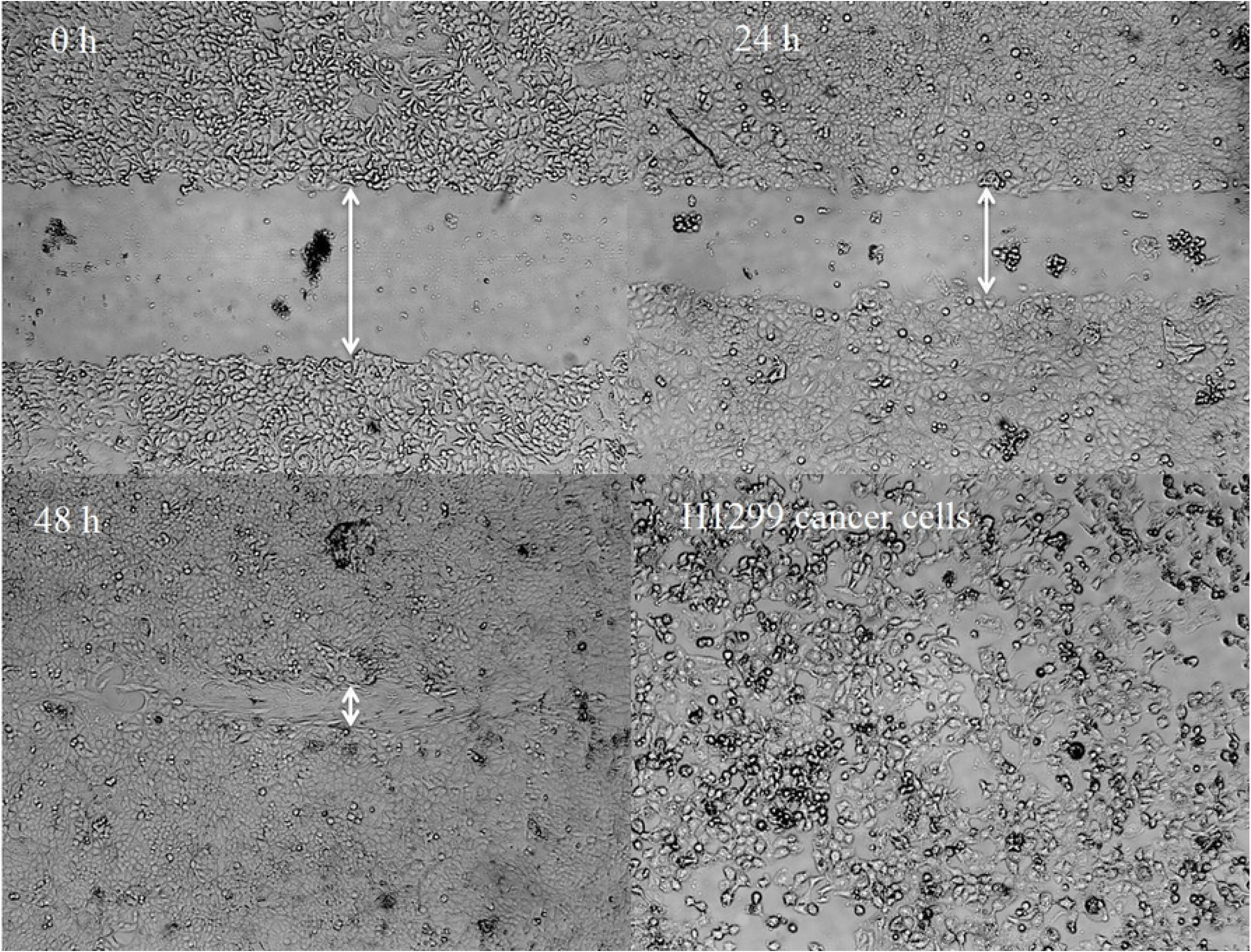
Effect of fertile HCF on H1299 human lung cancer cell migration. After 24 and 48 h, the scratch area was observed by an inverted microscope (A). The quantity of the migrated cells into the scratch area compared with the control group at 0, 24 and 48 h exposure times.

### 3.4. Assessment of mRNA expressions in Bax/Bcl-2 ratio, caspase-3, VGEF and Vimentin

Statistical results characterized that HCF-treated H1299 cells up-regulated Bcl-2 (an anti-apoptotic molecule) mRNA expression level compared to Bax (a pro-apoptotic molecule), suggesting less potent apoptosis of the lung cancer cell. As shown in Figure 4, the Bax/Bcl-2 ratio was decreased in HCF-treated H1299 cells (about 2-fold change) compared to control cells. The mRNA expression of Vimentin (as metastasis growth factor) and VGEF (Vascular endothelial growth factor indicating angiogenesis) genes in fertile HCF-treated H1299 cell line is illustrated in Figure 5. The findings indicated that after treatment of H1299 cell with fertile HCF, the mRNA expression levels of Vimentin (Fig. 5A), VGEF (Fig. 5B) were up-regulated to 1.7-fold change (*P*v; 0.0092) and 2.3-fold change (*P*v; 0.0033), respectively compared to control cell. Assessment of caspase-3 mRNA expression in HCF-treated H1299 cells was significantly decreased compared to control cells (*P*v; 0.0041; Fig. 6A).

**Figure 4:**
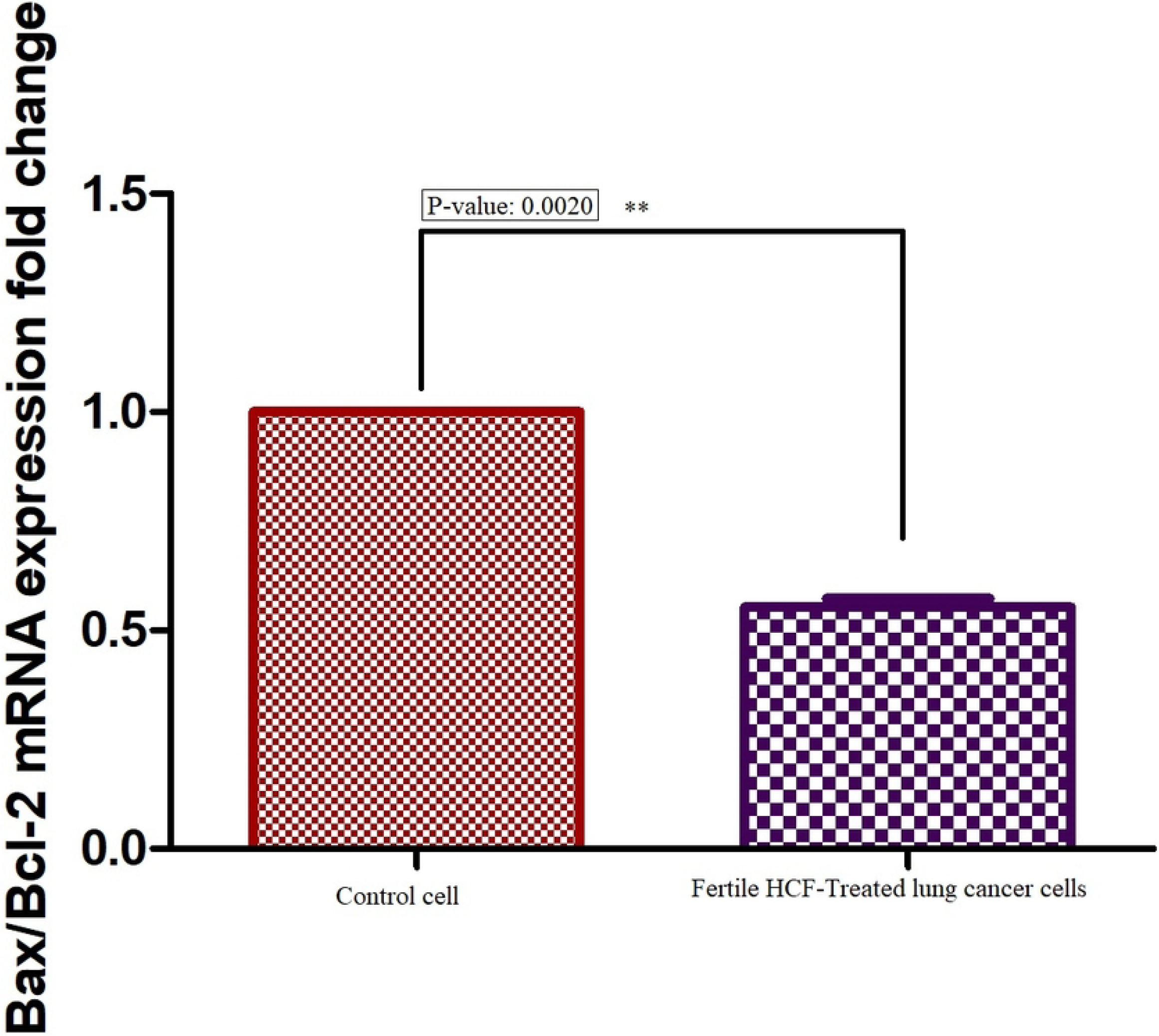
Bax/Bcl-2 mRNA expression ratio in fertile HCF-treated H1299 cell compared to control cells.

**Figure 5:**
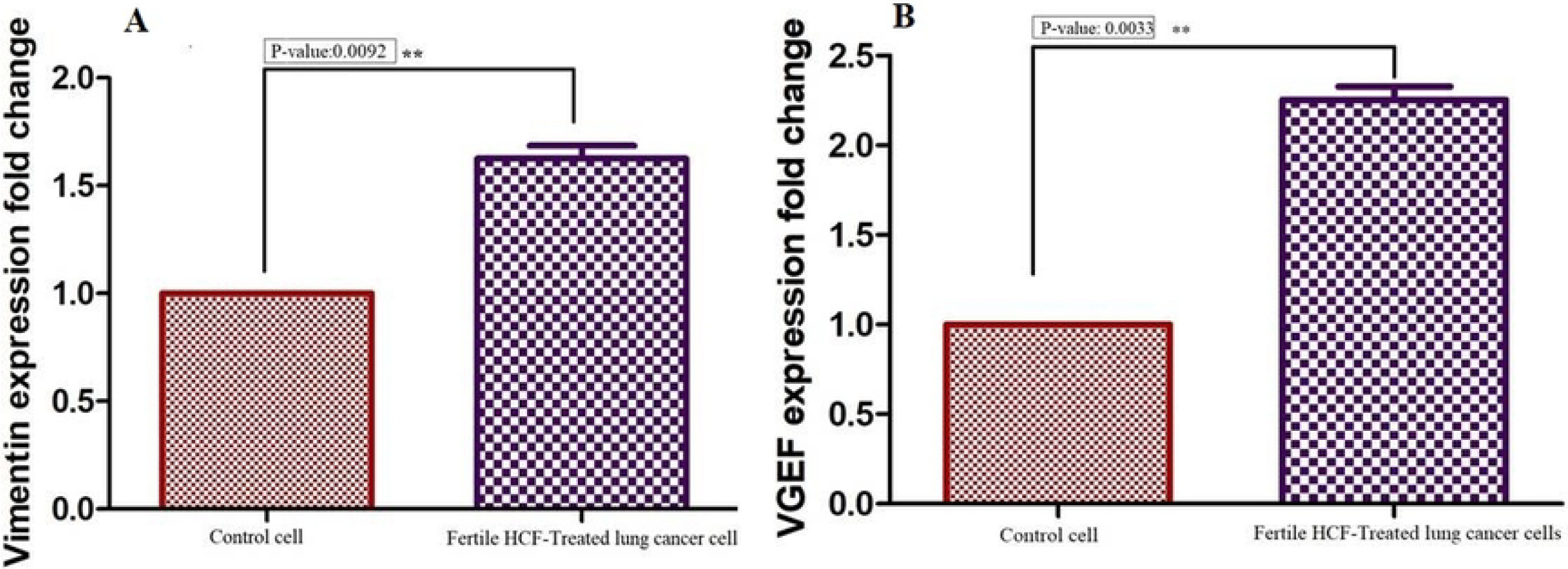
mRNA expression levels of Vimentin (**A**) and VGEF (**B**) in HCF-treated H1299 cell compared to control cell.

**Figure 6:**
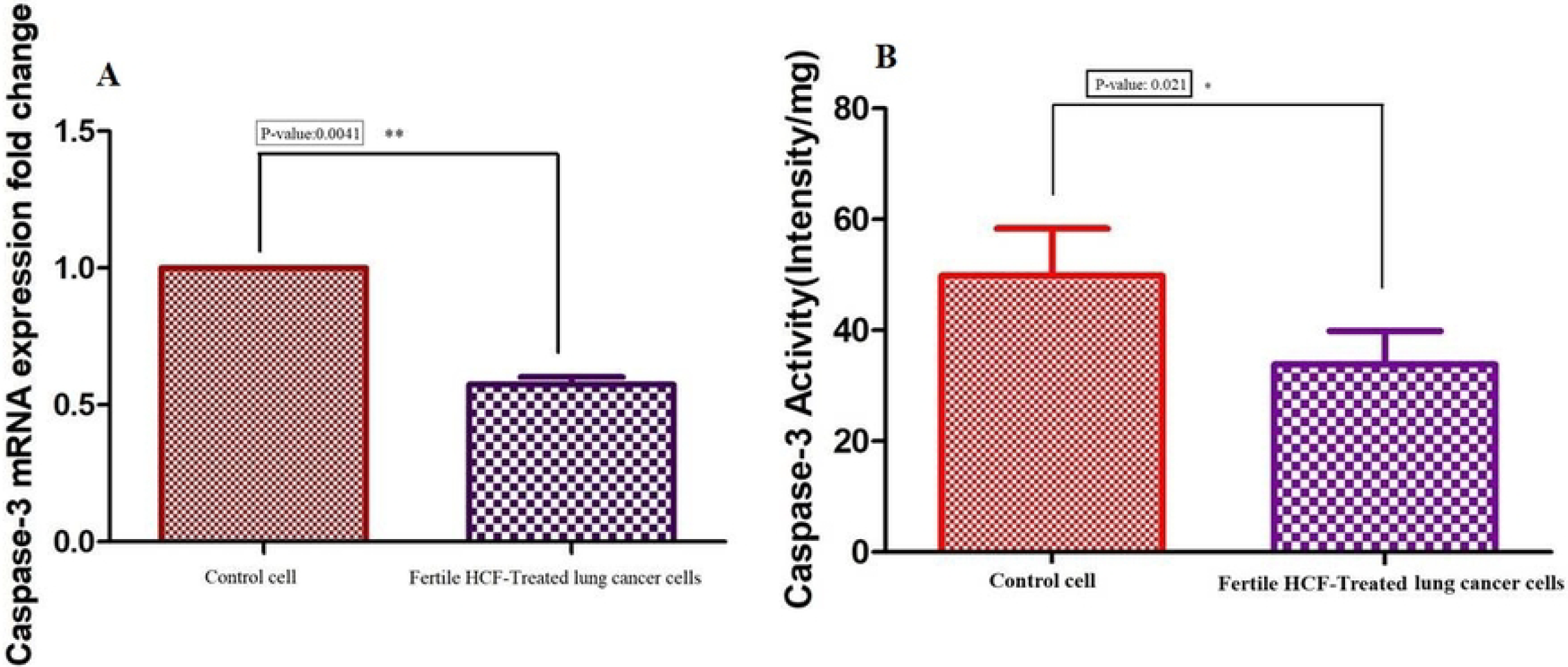
**A:** Evaluation of caspase-3 activity in cell extracts. Caspase-3 activity in fertile HCF-treated H1299 cell compared to control cell. **B**: mRNA expression level of caspase-3 in fertile HCF-treated H1299 cells compared to control cell.

### 3.5. Assessment of caspase-3 enzymatic activity

Results of fluorometric assay showed that caspase-3 enzymatic activity was meaningfully lower in HCF-treated H1299 cells than in control cells (*P*v; 0.021) (Fig. 6B).

## 4. Discussion

In this investigation, we evaluated the regulatory effects of lyophilized fertile HCF of *E. granulosus* s.s. on H1299 human lung cancer cells through decreasing apoptosis, increasing the cancer cells growth, migration of lung cancer cells as well as up-regulating VGEF and Vimentin genes.

The overall survival rate of lung cancer patients is unsatisfactory and most patients expire within a few months after diagnosis[18]. Therefore the importance of identification of predisposing factors in the progression and intensifying of lung cancer is essential, particularly in underlying/infectious diseases to choose well-suited treatments.

In terms of structural layers of pulmonary hydatid cyst, because of an absence of the adventitial outer layer and the thinness of the laminated layer, HCF antigens leakage into the thoracic space is feasible. According to the present results, in the immunological cross-talk between parasite and host, it can be inferred that any contamination of lung cancer patients with pulmonary hydatid cysts may intensify the severity of cancer through angiogenesis and tumorgenesis.

The anticancer property of HCF metabolites has been previously investigated in various cancer lines including breast, colon, melanoma, cervical, adenocarcinoma and pancreatic [5-7,19].

As well, Karadayi et al., (2013) have shown that collected sera from CE patients had a lethal impact on the growth of lung cancer cells *in vitro* [10]. Conversely, the present study indicated that the growth of lung cancer cells increases after exposure to lyophilized HCF of *E. granulosus* s.s. These discrepant findings may be justified by antigenic similarities/dissimilarities between parasite surface and host cancer cell, the type of cancer cell line, parasite ligand-host receptor interactions during invasion (docking process) and/or occurrence of resistant codons during the ectopic mutation [5,12].

Albeit, a dual function of miR-145 has been clarified as tumor suppressor and oncogene in cancers, however, the potential role of microRNAs especially miR-145 in the promotion/suppression of parasitic diseases has not been fully recognized yet [14]. Current findings indicate that miR-145 can enhance lung cancer cell growth after exposure to lyophilized hydatid cyst fluid of *E. granulosus* s.s. Therefore, using miR-145 replacement can exert a crucial role in reducing the migration and growth of lung cancer.

Hadighi et al., (2022) have shown that assessment of miR-145 in the plasma level of Iranian *Plasmodium vivax* malaria patients can introduce a promising diagnostic biomarker [20]. Farani et al., (2022) have also shown that over-expression of miR-145-5p and miR-146b-5p is related to decreased parasite burden in *Trypanosoma cruzi*-infected cardiomyoblasts which can candidate as promising biomarkers of parasite control to apply treatment methods in Chagas disease [21]. As well, Mohammadi et al., (2021) have shown that microRNA-365 enhanced apoptosis in A375 human melanoma cells line treated with fertile HCF [5]. Zhang et al., (2016) have demonstrated that HCF promotes peri-cystic fibrosis in CE through the increase in TβRII by suppressing miR-19 expression [22].

In this study, HCF-treated H1299 cancer cells up-regulated Bcl-2 mRNA expression level compared to Bax, suggesting a low apoptotic effect on the lung cancer cell. In consistent with the current study, Gao et al. (2018) have indicated that HCF up-regulated the procaspase-3 and Bcl-2 expression, and down-regulated the Bax expression, suggesting inhibition of melanoma cell apoptosis [19].

In conclusion, we down-regulated the miR-145 expression in H1299 human lung cancer cells, through treatment of lyophilized HCF antigens. Current results propose that induction of low expression of miR-145 in patients with hydatid cysts may be an oncogenic regulator of lung cancer growth through down regulation of apoptotic molecules and up regulation of angiogenesis and metastatic growth factor. Based on current findings, miR-145 may be a promising marker for the diagnosis of lung cancer patients co-infected with pulmonary hydatid cysts. To validate this assumption, further study is needed to assess microRNA profile and potent oncogenes *in vivo* setting.

## Acknowledgments

This is a report of a database from the thesis of Mr Hosein Mosajakhah registered in Tabriz University of Medical Sciences.

## Ethical approval

IR.TBZMED.VCR. REC.1401.095

